# Highly sensitive and multiplexed in situ protein profiling with cleavable fluorescent streptavidin

**DOI:** 10.1101/555615

**Authors:** Renjie Liao, Diego Mastroeni, Paul D. Coleman, Jia Guo

## Abstract

The ability to perform highly sensitive and multiplexed in situ protein analysis is crucial to advance our understanding of normal physiology and disease pathogenesis. To achieve this goal, here we develop an approach using cleavable biotin conjugated antibodies and cleavable fluorescent streptavidin (CFS). In this approach, protein targets are first recognized by the cleavable biotin labeled antibodies. Subsequently, CFS is applied to stain the protein targets. Though layer-by-layer signal amplification using cleavable biotin conjugated orthogonal antibodies and CSF, the protein detection sensitivity can be enhanced by at least 10 fold, compared with the existing methods. After imaging, the fluorophores and the biotins unbound to streptavidin are removed by chemical cleavage. The leftover streptavidin is blocked by biotin. Upon reiterative analysis cycles, a large number of different proteins with a wide range of expression levels can be unambiguously detected in individual cell in situ.

---

Comprehensive molecular profiling in single cells in situ holds great promise to reveal cell-to-cell variations and cell-microenvironment interactions, which are masked by population-based measurements^1,2^. Various methods^3–8^ have been developed for multiplexed single-cell analysis. An increasing number of studies have been focused on proteins, for its central roles in biological processes. Immunofluorescence (IF) is a well-established single-cell in situ protein analysis platform. However, on each specimen, only a couple of proteins can be profiled by IF, due to spectral overlap of commonly available organic fluorophores^9^.

To enable multiplexed in situ protein profiling, a number of methods^5–7,10–15^ have been developed recently. In these methods, the detection tags are either conjugated to the primary antibodies or secondary antibodies. Without signal amplification, the existing methods have limited detection sensitivity, which impede the analysis of low expression proteins. Additionally, due to the low sensitivity of the current methods, long imaging exposure time is required, which results in limited sample throughput and long assay time.

Here, we report a highly sensitive and multiplexed in situ protein analysis approach. In this method (Figure 1), reiterative protein staining is applied to achieve comprehensive single-cell protein analysis. Each staining cycle is composed of five major steps. First, proteins of interest are targeted by cleavable biotin labeled primary antibodies and cleavable fluorescent streptavidin (CFS). Second, the specimen is incubated with a cleavable biotin labeled orthogonal antibody followed by CFS. This layer-by-layer signal amplification step can be repeated several times to achieve the desired signal intensities. Third, the specimen is imaged to generate quantitative single-cell protein expression profiles. Fourth, the fluorophores and the biotins unbound to streptavidin are efficiently removed by chemical cleavage. Finally, the leftover streptavidin is blocked with biotin. Through reiterative cycles of staining, amplification, imaging, cleavage and streptavidin blocking, a large number of proteins expressed with a wide range of expression levels can be characterized in single cells in situ.

**Figure 1.**
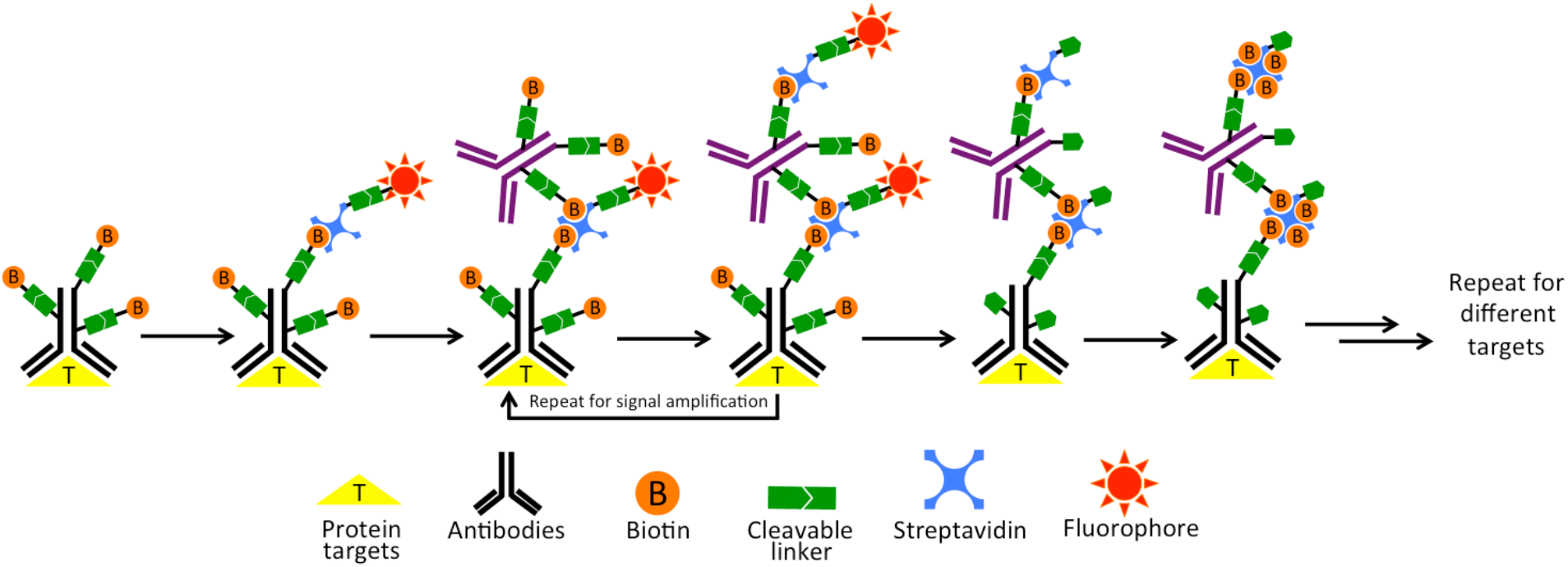
Highly sensitive and multiplexed in situ protein profiling with cleavable fluorescent streptavidin (CFS). In each cycle, the protein of interest is first targeted by cleavable biotin labeled primary antibody, and then stained with CFS. Though layer-by-layer signal amplification using cleavable biotin conjugated antibodies and CFS, highly sensitive protein detection is achieved. After imaging, the fluorophores and the biotins unbound to streptavidin are chemically cleaved and subsequently streptavidin is blocked by biotin. Through reiterative cycles of target staining, signal amplification, fluorescence imaging, chemical cleavage and streptavidin blocking, comprehensive protein profiling can be achieved

To demonstrate the feasibility of this approach, we conjugated biotin to antibodies through a disulfide bond based cleavable linker and Cy5 to streptavidin through an azide based cleavable linker, according to previously describe method^12^. In this way, both biotin and Cy5 can be simultaneously removed by the reducing reagent tris(2-carboxyethyl)-phosphine (TCEP).

We then evaluated the detection sensitivity of our approach by comparing it with direct and indirect immunofluorescence. Protein Ki67 in Hela cells was stained with these 3 methods with the same concentration of primary antibodies (Figure S1A). Without any signal amplification steps, the CFS method is ∼4.5 times more sensitive than direct immunofluorescence without losing the staining resolution. The detection sensitivity of the CFS method is comparable to that of indirect immunofluorescence (Figure S1B). With 4 rounds of signal amplification, the original staining intensities were increased by more than 10 times (Figure 2). These results demonstrate that our approach enhances the detection sensitivity of the existing methods by at least one order of magnitude.

**Figure 2.**
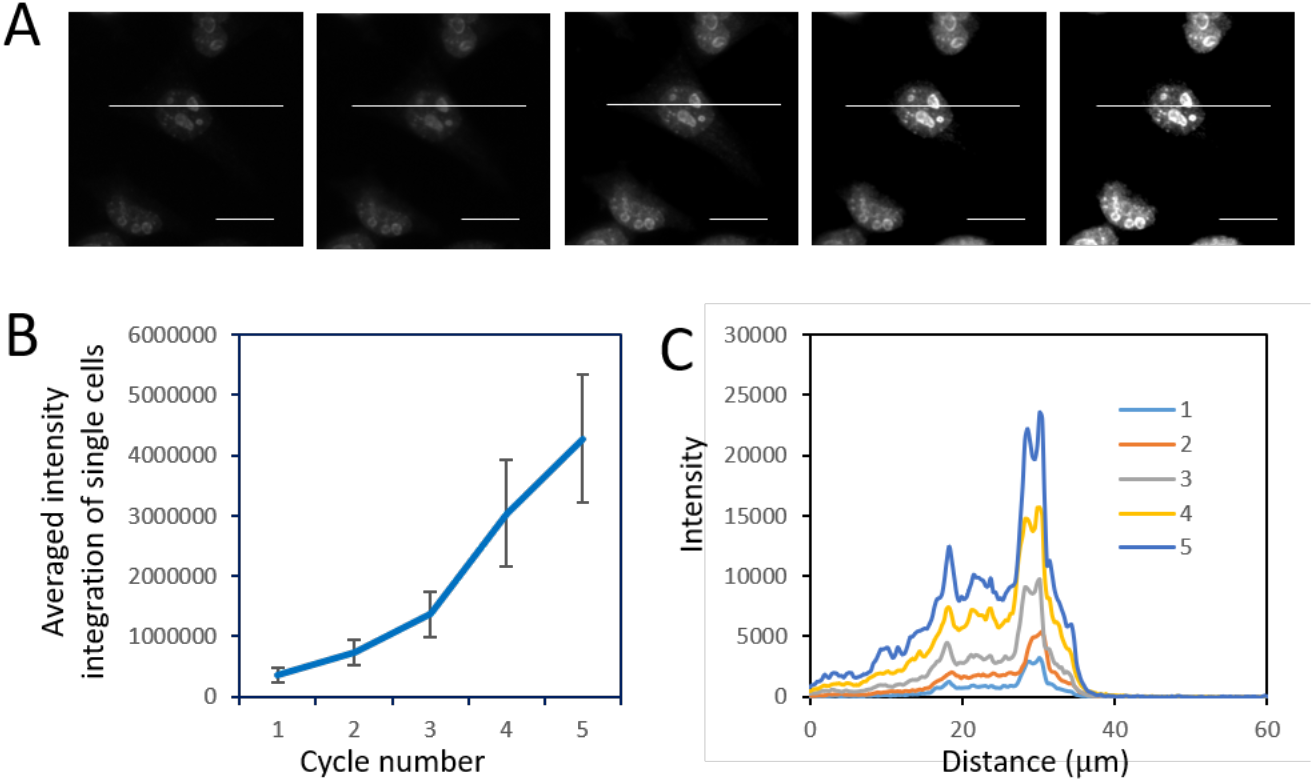
A) Fluorescent images of protein Ki67 stained with 1 to 5 amplification cycles in HeLa cells. B) Averaged signal integration in single cells (n=30) in amplification cycles 1 to 5. Error bars, standard deviation. C) Fluorescence intensity profiles corresponding to the indicated line positions in amplification cycles 1 to 5. Scale bars, 20 μm.

To enable multiplexed protein analysis by reiterative analysis cycles, three major requirements exist. (1) Fluorescence signals need to be efficiently erased by chemical cleavage. (2) The biotins that are not bounded to streptavidin have to be efficiently removed to avoid false positive signals in the next staining cycle. (3) Streptavidin needs to be efficiently blocked before the next staining cycle. To assess whether these three requirements are met by the CFS approach, we stained protein Ki67 with 1 to 5 amplification cycles in HeLa cells, and first quantified the cleavage efficiency (Figure S2). After TCEP incubation, ∼95% of signal was removed regardless of the number of the amplification rounds. To test whether the biotins unbounded to streptavidin can be removed by TCEP, we stained protein Ki67 with 1 to 5 amplification cycles (Figure S3). After TCEP cleavage, the cells were incubated the CFS, again. No further fluorescence signal enhancement was introduced, suggesting that the free biotins are efficiently removed during the cleavage step. To evaluate the streptavidin blocking efficiency, we stained protein Ki67 with 1 to 5 amplification cycles (Figure S4). Subsequently, the cells were incubated with TCEP and then with biotin to block streptavidin. Another round of signal amplification was applied and no further fluorescence signal enhancement was detected. These results indicate that streptavidin is efficiently blocked by biotin.

Archived tissues are important biological samples to study normal physiology and disease pathogenesis. Formalin fixed, paraffin embedded (FFPE) tissue is the most common form of archived tissues in clinics and pathology labs^20^. FFPE tissues often bear high autofluorescence^11^ and the partially degraded proteins^21^, which makes them difficult to be profiled by the fluorescence imaging methods with low detection sensitivities. To demonstrate the feasibility of applying the CFS approach to analyze FFPE tissues, we stained H3K4me3 in a FFPE human brain tissue (Figure 3). With 2 rounds of signal amplification, the signal-to-background ratio was significantly improved. After cleavage, the fluorescence signal was efficiently removed. Another round of signal amplification cycle after cleavage and streptavidin blocking did not further increase the staining intensities. These results confirm that the flurophores and the free biotins can be efficiently removed by TCEP and streptavidin can be efficiently blocked by biotin. These results also imply that the CFS approach can be successfully applied to quantify the partially degraded proteins in highly fluorescent FFPE tissues.

**Figure 3.**
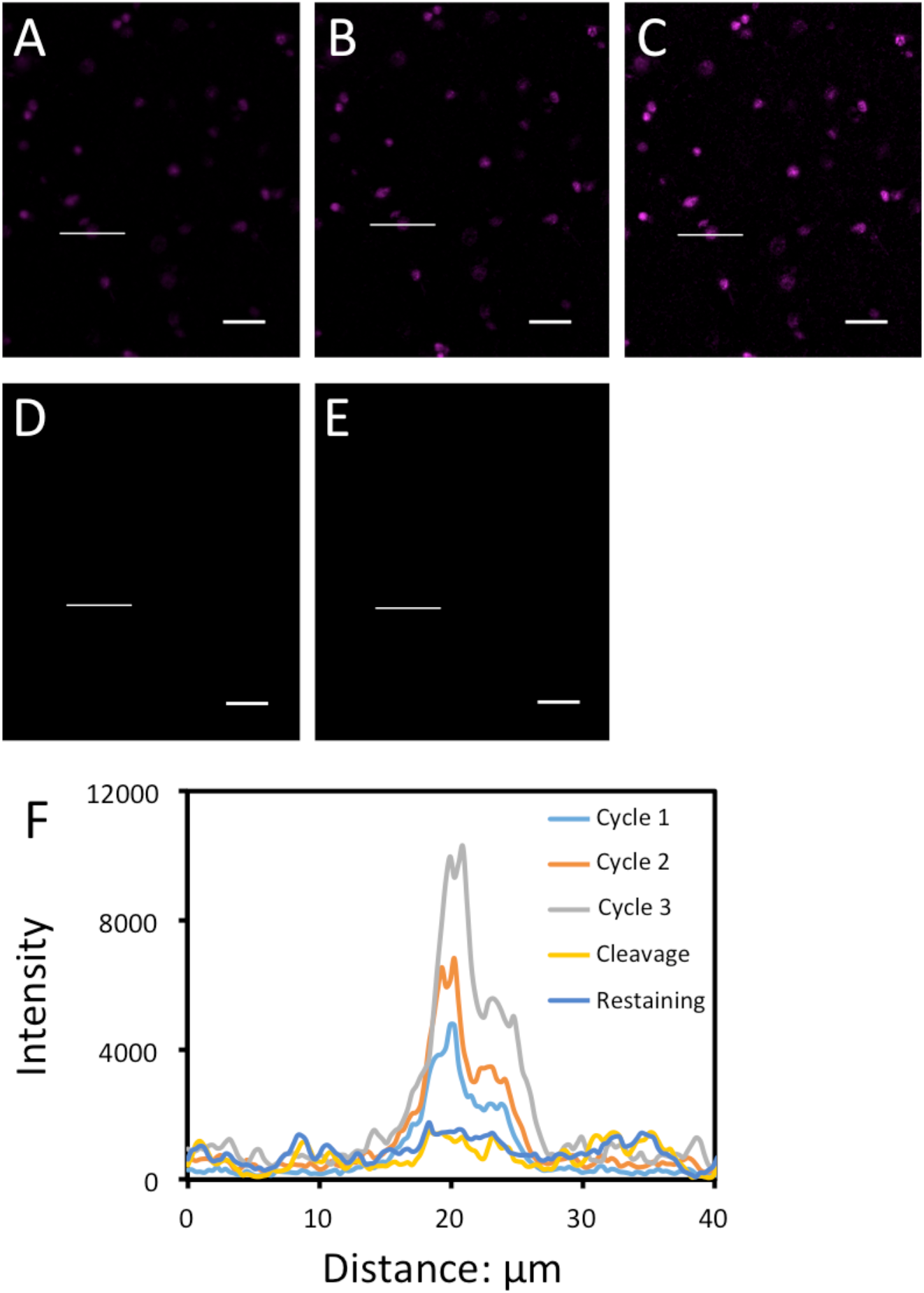
Protein H3K4me3 in a human FFPE brain tissue was stained with CFS in amplification cycles A) 1, B) 2 and C) 3. D) Afterwards, the stained tissue was incubated with TCEP. E) Following chemical cleavage and streptavidin blocking, the tissue was incubated with cleavable biotin conjugated antibodies and CFS again. F) Fluorescence intensity profiles corresponding to the indicated line positions in (A) to (E). Scale bars, 25 μm.

In summary, we have designed and synthesized CFS and demonstrated this multiplexed in situ protein analysis approach enhances the detection sensitivity of the existing approaches by at least one order of magnitude. With the dramatically improved sensitivity, this approach enables the quantitative analysis of low expression proteins, especially in the highly autofluorescent tissue samples. The multiplexing capacity of this approach depends on two factors: the number of reiterative analysis cycles and the number of proteins quantified in each cycle. We have shown previously that the protein antigenicity is preserved after the incubation with TCEP for 24 hours^12^, which suggests that more than 40 cycles can be carried out on the same specimen. In each cycle, varied protein targets can be first recognized by primary antibodies labeled with distinct cleavable haptens, such as biotin, fluorescein, TAMRA, and digoxigenin (DIG). Subsequently, streptavidin, anti-fluorescein, anti-TAMRA, and anti-DIG antibodies labeled with different fluorophores can be applied to stain the protein targets and amplify the signals. In this way, at least four proteins can be quantified simultaneously in each cycle. Thus, we anticipate this method has the potential to analyze >100 protein targets in the same specimen. This highly multiplexed and sensitive in situ protein profiling technology will have wide applications in systems biology and biomedical research.

## Acknowledgments

This work is supported by the National Institute of General Medical Sciences (1R01GM127633), the National Institute Of Allergy And Infectious Diseases (R21AI132840), Arizona State University startup funds, and Arizona State University/Mayo Clinic seed grant (ARI-219693).

