## Supplementary material for "Highly sensitive and multiplexed in situ protein profiling with cleavable fluorescent streptavidin": SI

Table of Contents

1. General information……………………………………………………………………………………S3

2. Protein staining with cleavable fluorescent streptavidin (CFS) in cells……………………S3

3. Quantification of the fluorophore cleavage efficiency…………………………………………S5

4. Quantification of the biotin cleavage efficiency……………………………………………………S5

5. Quantification of the streptavidin blocking efficiency…………………………………………S5

6. Protein staining in a brain FFPE tissue……………………………………………………………S5

7. Imaging and data analysis……………………………………………………………………………S6

8. Figure S1……………………………………………………………………………………………………S7

9. Figure S2……………………………………………………………………………………………………S8

10. Figure S3…………………………………………………………………………………………………S9

11. Figure S4…………………………………………………………………………………………………S10

### 1. General information

Chemicals and solvents were purchased from Sigma-Aldrich or TCI America, used directly without further purification, unless otherwise noted. Bioreagents were purchased from Invitrogen, unless otherwise indicated.

### 2. Protein staining with cleavable fluorescent streptavidin (CFS) in cells

#### Cell culture

Hela CCL-2 cells (ATCC) were maintained in Dulbelcco's modified Eagle's Medium (DMEM) supplemented with 10% fetal bovine serum, 100 U/mL penicillin and 100 g/mL streptomycin in a humidified atmosphere at 37 $^{o}$C with 5% $CO_2$. Cells were plated on chambered coverglass (0.2 ml medium/chamber) (Thermo Fisher Scientific) and allowed to reach 60% confluency in 1-2 days.

#### Cell fixation and permeabilization

Cultured Hela CCL-2 cells were fixed with 4% formaldehyde (Polysciences) in 1X PBS at 37 °C for 15 min, followed by washing with 1X PBS, 3 x 5 min. Cells were then permeabilized with PBT (0.1% Triton-X 100 in 1X PBS) for 10 min at room temperature, and subsequently washed 3 times with 1X PBS, each for 5 min.

#### Preparation of cleavable fluorescent streptavidin

Cleavable Cy5 NHS ester was prepared according to the literature[1]. To 20 μL of streptavidin solution at a concentration of 1 mg/mL, 1 nmol of cleavable Cy5 NHS ester and 2 μL of $NaHCO_3$ solution (1 M) was added. The mixture was incubated in dark and room temperature for 15 min. The labeled streptavidin was purified by p-6 biogel column.

#### Preparation of Biotin-SS-Ab

To 20 μL of primary antibody solution at a concentration of 1 mg/mL, 3 nmol of EZ link Sulfo-NHS-SS-Biotin (Thermo Fisher Scientific) and 2 μL of $NaHCO_3$ solution (1 M) was added. The mixture was incubated in dark and room temperature for 15 min, and then the conjugation product was purified by p-6 biogel column.

**Immunofluorescence with CFS**

Fixed Hela cells were incubated with antibody blocking buffer (10% normal goat serum (v/v), 1% bovine serum albumin (w/v), 0.1 Triton-X 100 in 1X PBS) for 1 h at room temperature, and then washed 3 times with PBT, each for 5 min. To block the cell endogenous biotin, the cells were treated with 0.1 mg/mL streptavidin in 1X PBS for 15 min at room temperature, and washed 3 times with 1X PBS, each for 5 min. Subsequently, the cells were incubated with 0.5 mg/mL biotin in 1X PBS for 30 min at room temperature, and washed with 1X PBS 3 times, each for 5 min. After blocking, the cells were incubated with Biotin-SS-Ab in antibody blocking buffer (concentration varies and suggested by manufacturers) for 45 min at room temperature, and washed with PBT for 3 times, each for 10 min. Subsequently, the cells were incubated with 10 ng/μL cleavable fluorescent streptavidin in 1% BSA in PBT for 30 min, and washed 3 times with 1X PBS, each for 5 min. The cells were washed with GLOX buffer (0.4% glucose and 10 mM Tris HCl in 2 X SSC) for 1-2 min at room temperature, and then imaged in GLOX solution (0.37 mg $mL^{-1}$ glucose oxidase and 1% catalase in GLOX buffer).

**Amplification**

To amplify the staining signal, the cells were incubated with cleavable biotin conjugated orthogonal secondary antibodies in 1% BSA in PBT at a concentration of 10 ng/μL for 30 min, and then washed 3 times with 1X PBS, each for 5 min. Afterwards, the cells were incubated with cleavable fluorescent streptavidin in 1% BSA in PBT at a concentration of 10 ng/μL, and again washed 3 times with 1X PBS, each for 5 min. Multiple amplification cycles can be repeated to obtain the desired signal intensity.

**Cleavage**

Cleavage was performed by incubating the specimen with tris(2-carboxyethyl)phosphine (TCEP, pH=9.5, 100 mM in deionized water) for 30 min at 37 $^{o}$C. Subsequently, the cells were washed 3 times with PBT and 3 times with 1X PBS, each for 5 min.

**Blocking**

After cleavage, the cells were incubated with 0.5 mg/mL biotin in 1X PBS for 30 min at room temperature, and then washed 3 times with 1X PBS, each for 5 min.

### 3. Quantification of the fluorophore cleavage efficiency

Fixed and blocked Hela CCL-2 cells were incubated with 0.01 mg/mL cleavable biotin labeled rabbit anti-Ki67 (Thermo Fisher Scientific) for 45 min. Subsequently, the cells were stained by 0.01 mg/mL cleavable fluorescent streptavidin. Then, one, two, three and four rounds of amplification were applied to different sets of cells. In each round of amplification, the cells were first incubated with cleavable biotin labeled goat-anti-chicken antibodies and then with cleavable fluorescent streptavidin. The cells were then incubated with TCEP (100 mM, pH=9.5) for 30 min at 37 $^{o}$C. Subsequently, the cells were washed 3 times with PBT and 3 times with 1X PBS, each for 5 min.

### 4. Quantification of the biotin cleavage efficiency

Fixed and blocked Hela CCL-2 cells were incubated with 0.01 mg/mL cleavable biotin labeled rabbit anti-Ki67 (Thermo Fisher Scientific) for 45 min. Subsequently, cells were stained by 0.01 mg/mL cleavable fluorescent streptavidin. Following that, one, two, three and four rounds of amplification were applied to different sets of cells. In each round of amplification, the cells were first incubated with cleavable biotin labeled goat-anti-chicken antibodies and then with cleavable fluorescent streptavidin. Biotin and fluorophores were cleaved by TCEP (100 mM, pH=9.5). The cells were then incubated with cleavable fluorescent streptavidin.

### 5. Quantification of the streptavidin blocking efficiency

Fixed and blocked Hela CCL-2 cells were incubated with 0.01 mg/mL Sulfo-NHS-SS-Biotin labeled rabbit anti-Ki67 (Thermo Fisher Scientific) for 45 min. Subsequently, cells were stained by 0.01 mg/mL cleavable fluorescent streptavidin. Following that, one, two, three and four rounds of amplification were applied to different sets of cells. In each round of amplification, the cells were first incubated with cleavable biotin labeled goat-anti-chicken antibodies and then with cleavable fluorescent streptavidin. Biotin and fluorophores were cleaved by TCEP (100 mM, pH=9.5). The cells were blocked with 0.5 mg/mL biotin. The cells were incubated with cleavable biotin labeled goat anti-chicken and then cleavable fluorescent streptavidin.

### 6. Protein staining in a brain FFPE tissue

#### Deparaffinization and antigen retrieval

A brain FFPE tissue slide was deparaffinized in xylene for 3 times, 10 min for each. Then the slide was immersed in 100% ethanol for 2 min, 95% ethanol for 1 min, 70% ethanol for 1 min, 50% ethanol for 1 min, 30% ethanol for 1 min. The slide was rinsed with deionized water.

A combination of 'heat induced antigen retrieval' (HIAR) and 'enzymatic antigen retrieval' was used. HIAR was done using a pressure cooker (Cuisinart). The slide was immersed in antigen retrieval buffer (10 mM sodium citrate, 0.05% Tween 20, pH=6.0), and water-bathed in pressure cooker for 20 min with the 'High pressure' setting. Subsequently, the slide was rinsed 3 times with 1X PBS, each for 5 min. The slide were treated with pepsin digest-all 3 (Life Technologies) for 10 min, and then washed 3 times with 1X PBS, each for 5 min.

#### Protein staining in FFPE tissues

To block the endogenous biotin, the slide was treated with 0.1 mg/mL streptavidin in 1X PBS for 15 min at room temperature, and washed 3 times with 1X PBS, each for 5 min. Subsequently, the slide were incubated with 0.5 mg/mL biotin in 1XPBS for 30 min at room temperature, and washed 3 times with 1X PBS, each for 5 min. The slide was incubated with 0.01 mg/mL cleavable biotin labeled rabbit anti-H3K4me3 (Cells Signaling) in antibody blocking buffer for 45 min, and washed 3 times with PBT, each for 10 min. The slide was stained by 0.01 mg/mL cleavable fluorescent streptavidin for 30 min, and then washed 3 times with 1X PBS, each for 5 min. Two cycles of amplification were applied. In each round of amplification, the cells were first incubated with cleavable biotin labeled goat-anti-chicken antibodies and then with cleavable fluorescent streptavidin. After imaging, the slide was incubated with TCEP (100 mM, pH=9.5) for 30 min at 37 $^{o}$C, and washed 3 times with PBT and 3 times with 1X PBS, each for 5 min. Streptavidin was blocked with 0.5 mg/mL Biotin. The tissue was restained with 0.01 mg/mL cleavable biotin labeled goat anti-chicken and then 0.01 mg/mL cleavable fluorescent streptavidin.

### 7. Imaging and data analysis

Stained cells and brain FFPE tissue were imaged under a Nikon Ti-E epifluorescence microscope equipped with 20x objective. Images were captured using a CoolSNAP HQ2 camera and Chroma filter 49009. Image data was analyzed with NIS-Elements Imaging software.

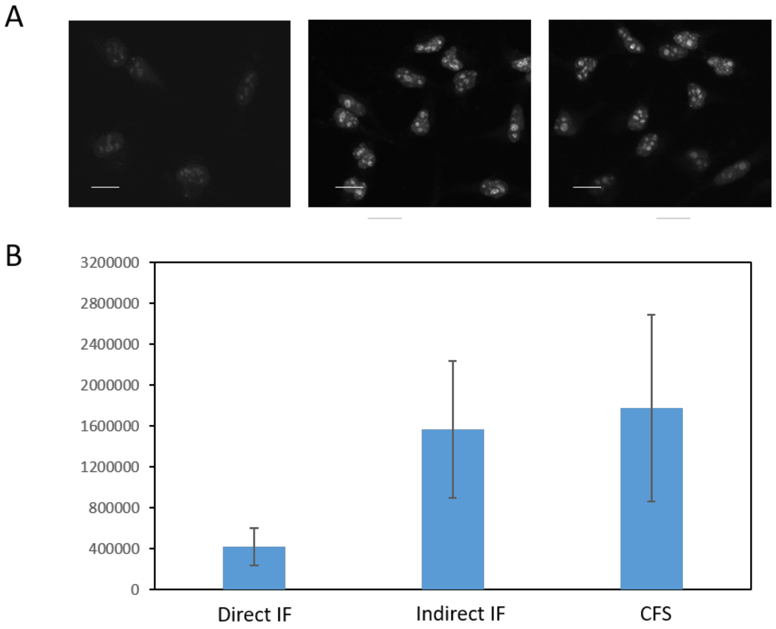


**Figure S1**. A) Fluorescent images of protein Ki67 stained with direct IF (left), indirect IF (middle) and cleavable fluorescent streptavidin (CFS). B) Comparison of normalized staining intensity (n=30) for 3 methods. Scale bars, 20 μm.

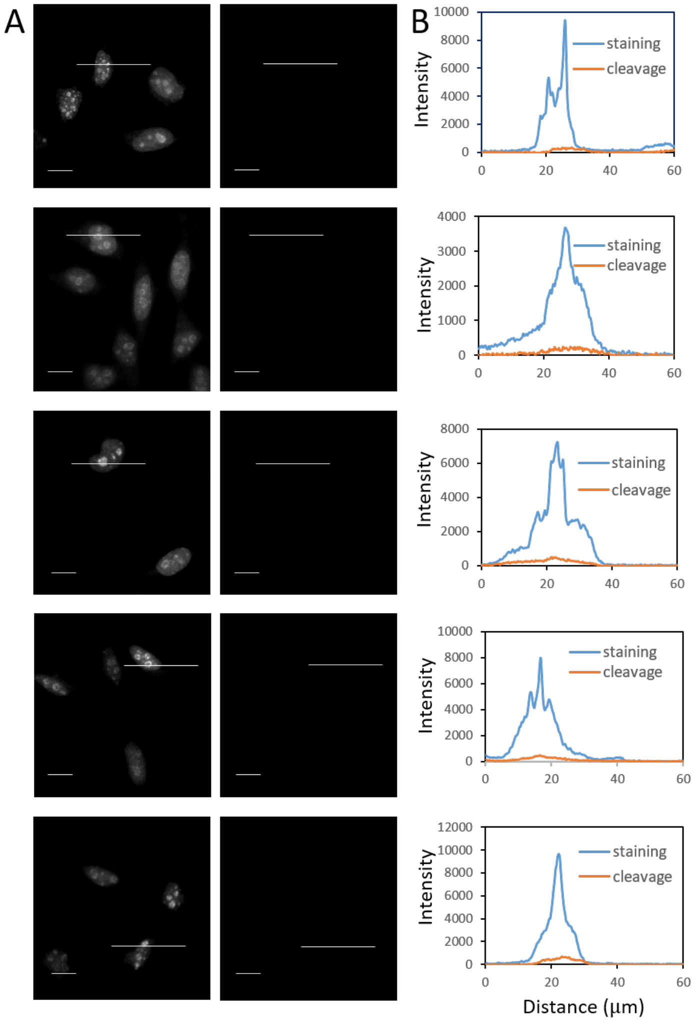


**Figure S2**. A) Fluorescent images of protein Ki67 stained with 1 to 5 amplification cycles in Hela cells (left column) and those after cleavage (right column). The exposure time in amplification cycles 1 to 5 is 1 s, 500 ms, 250 ms, 125 ms, 62 ms, respectively. (B) Fluorescence intensity profiles corresponding to the indicated line positions in amplification cycles 1 to 5. Scale bars, 20 μm.

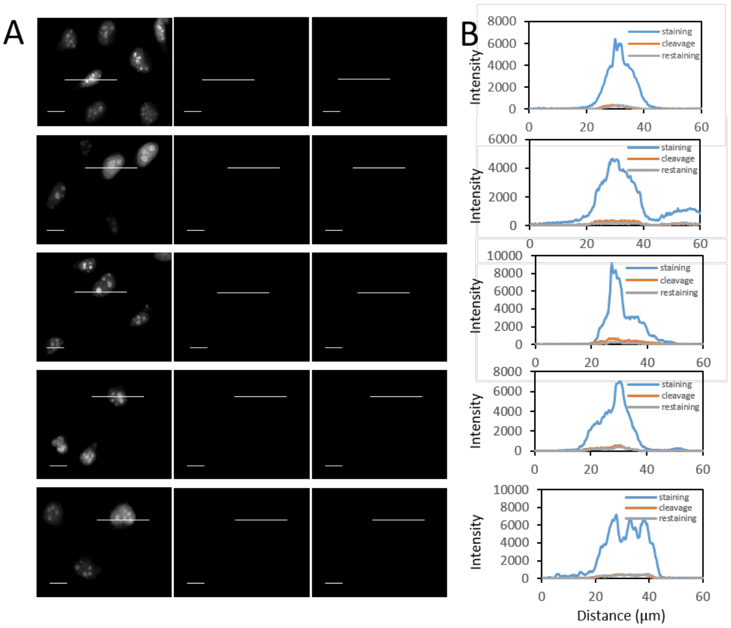


**Figure S3.** A) Fluorescent images of protein Ki67 stained with 1 to 5 amplification cycles in HeLa cells (left column) and after cleavage (middle column) and restained with CFS (right column). The exposure time in amplification cycles 1 to 5 is 1 s, 500 ms, 250 ms, 125 ms, 62 ms, respectively. (B) Fluorescence intensity profiles corresponding to the indicated line positions in amplification cycles 1 to 5. Scale bars, 20 μm.

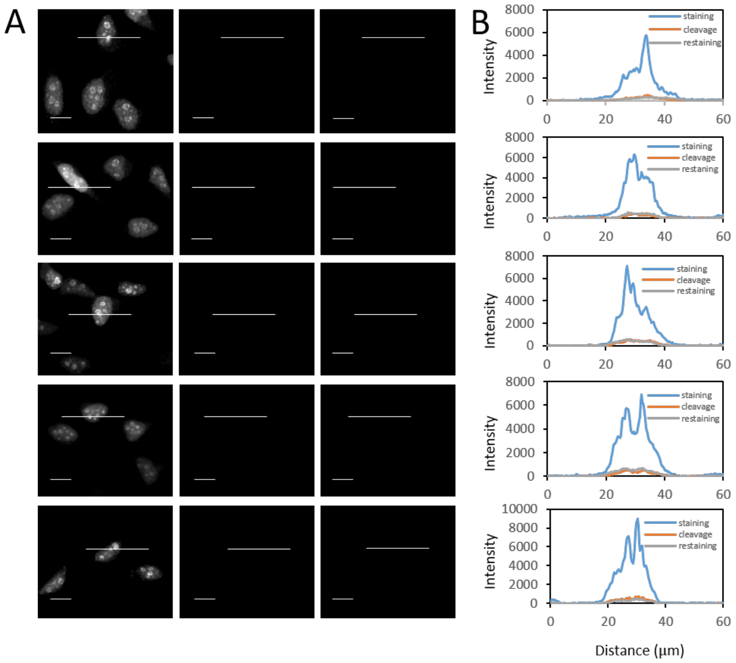


**Figure S4**. A) Fluorescent images of protein Ki67 stained with 1 to 5 amplification cycles in HeLa cells (left column) and those after cleavage (middle column). Following streptavidin blocking, the cells were restained with cleavable biotin labeled antibodies and CFS (right column). The exposure time in amplification cycles 1 to 5 is 1 s, 500 ms, 250 ms, 125 ms, 62 ms, respectively. B) Fluorescence intensity profiles corresponding to the indicated line positions in amplification cycles 1 to 5. Scale bars, 20 μm.
